# Knockdown of the fly spliceosome component *Rbp1*(orthologue of *SRSF1*) extends lifespan

**DOI:** 10.1101/2025.09.18.677196

**Authors:** Dan J. Hayman, Mirre J. P. Simons

## Abstract

Biological regulation is a highly intricate process and involves many layers of complexity even at the RNA level. Alternative splicing is crucial in the regulation of which components of a protein-coding gene are spliced into a translatable mRNA. During ageing splicing becomes dysregulated and alternative splicing has been shown to be involved in disease and known anti-aging treatments such as dietary restriction (DR) and mTOR suppression. In prior work we have shown that DR and mTOR suppression modulate the expression of the spliceosome in the fly (*Drosophila melanogaster*). Here, we manipulated the five top genes that change in expression in both these treatments. We found that knockdown (using conditional *in vivo* RNAi in adults) of some spliceosome components rapidly induce mortality, whereas one, *Rbp1*, extends lifespan. Treatments that have more instant benefits on longevity are more translatable. We therefore subsequently repeated the Rbp1 experiment, but initiating *Rbp1* at later stages in adult life. We find that irrespective of age of induction, knockdown of *Rbp1* extends lifespan. Our results posit the spliceosome itself as a hub of regulation that when targeted can extend lifespan, rendering it a promising target for geroscience.

## Introduction

Ageing is characterised by a diverse range of molecular and cellular alterations and is the strongest risk factor for all age-related diseases (Guo et al., 2022; López-Otín et al., 2023). The geroscience hypothesis therefore states that if we can target and treat ageing we will prevent all major debilitating age-related diseases (Kennedy et al., 2014). The two best studied treatments which positively impact health and lifespan across species are restriction (DR) and suppression of mammalian target of rapamycin (mTOR) (Garratt et al., 2016; Green et al., 2022; Papadopoli et al., 2019; Simons et al., 2013). Dietary restriction is conserved across evolution, spanning from yeast to mammals (Fontana et al., 2010; Gautrey et al., 2025). It remains unclear, however, which precise nutrients enable the beneficial effects of DR and what the exact downstream mechanisms of DR are (Gautrey & Simons, 2022; Green et al., 2022; Selman, 2014). The beneficial lifespan extending effects of suppression of mTOR appear to similarly be conserved across species (Garratt et al., 2016), but downstream mechanisms again are similar to DR not fully elucidated and distributed across many different physiological and molecular pathways. For example, mTOR expression has been reported to both increase and decrease with age, depending on, sex, tissue, and other specific conditions (Baar et al., 2016; Chen et al., 2009; Papadopoli et al., 2019). A shared feature of both mTOR suppression and DR is however that they both exert widespread effects on alternative splicing irrespective of species (Heintz et al., 2017; Rhoads et al., 2018; Simons et al., 2019; Tabrez et al., 2017). Both mTOR suppression and DR may therefore orchestrate physiology that promotes healthy ageing through changes in genome-wide splicing.

Alternative splicing is the process regulating which transcribed components of a protein-coding gene are spliced into a translatable mRNA determining a large proportion of the complexity of the proteome (Marasco & Kornblihtt, 2023; Modrek & Lee, 2002). Splicing changes rapidly in response to the environment (Singh & Ahi, 2022) and patterns of splicing are heritable (Kwan et al., 2007). Alternative splicing is a major determinant of organismal complexity and abnormal splicing events are implicated in disease and ageing (Bhadra et al., 2020; Marasco & Kornblihtt, 2023). Indeed, a genome wide dysregulation of alternative splicing is observed during ageing (Holly et al., 2013; Latorre & Harries, 2017; Li et al., 2017). The molecular machinery through which splicing is carried out and regulated is known as the spliceosome; a large dynamic complex of approximately 100 different proteins as well as snRNPs and small nuclear RNA (snRNA). Several other proteins trigger the assembly of the spliceosome, even if they are not themselves part of the spliceosome “core” (Kastner et al., 2019; Wilkinson et al., 2020).

Each spliceosome component is recruited to the spliceosome as the complex forms around a 5’ splice site, and each mediates specific parts of the splicing reaction (Plaschka et al., 2019; Yan et al., 2019). As each component is recruited (some individually, others as part of a complex), the spliceosome changes conformation and progresses through different stages of the splicing reaction (Wilkinson et al., 2020). Each component therefore has an important, albeit in some cases small, role to play in the splicing of mRNA. Previous studies have manipulated the expression of individual spliceosome genes and identified, importantly but also perhaps unsurprisingly, wide-reaching splicing and spliceosome regulatory changes (Rogalska et al., 2024). Intriguingly, the manipulation of a single individual spliceosome component can modulate lifespan; overexpression of one spliceosome component gene in *C. elegans, sfa-1*, extends lifespan, whereas knockdown of *sfa-1* negates the pro-longevity phenotypes of mTOR suppression and DR (Heintz et al., 2017). Modulating the spliceosome may thus have the potential to mimic the health benefits of both DR and mTOR suppression.

Here, we manipulated the spliceosome components that show the strongest transcriptomic changes in response to the rejuvenating effects of early-life mTOR suppression and DR in flies. Three spliceosome components truncated lifespan substantially when knocked down (*Sf3b1, barc* and *Prp5*), showing they are essential for life. One splicing factor, *Rbp1*, increases lifespan when knocked down, and similarly when this knockdown was induced later in life. Modulation of the spliceosome therefore holds promise to achieve pro-longevity effects. As such the spliceosome provides a model to distill how pro-longevity effects are orchestrated on a whole organism level.

## Results

### The expression of spliceosome genes change consistently between DR and transient mTOR suppression

We tested using previously generated transcriptomes in our group whether the spliceosome changed concordantly across treatments. Suppression of mTOR in early adult life (using RNAi) has long lasting benefits which we have suggested previously are mediated via the spliceosome (Simons et al., 2019). We combined this dataset with a transcriptome measuring the response to DR within 48 hours. As the fly responds rapidly to DR in terms of a reduction in mortality rate (Good & Tatar, 2001; Mair et al., 2003; McCracken et al., 2020; Whitaker et al., 2014) changes occurring in the transcriptome during this time are devoid of longer term compensation that will not be causal to the extension in lifespan observed. Previous work used microarrays (Whitaker et al., 2014), we recently generated next generation sequencing (using the setup described in Charles, 2022; Gautrey et al., 2025; McCracken et al., 2020). We compared whether the same spliceosome or spliceosome-regulatory genes (identified as genes which were in Gene Ontology terms featuring the word “spliceosome”; see Table S1) were differentially expressed in both the mTOR and DR datasets, and whether they were modulated with concordant directionality. Of the 280 total genes in the spliceosome-related Gene Ontology terms, 165 genes were significantly differentially expressed upon mTOR suppression and 36 upon DR. 32 of these genes were differentially expressed with the same directionality in both DR and transient mTOR suppressed conditions (Figure 1 and Table S2).

**Figure 1.**
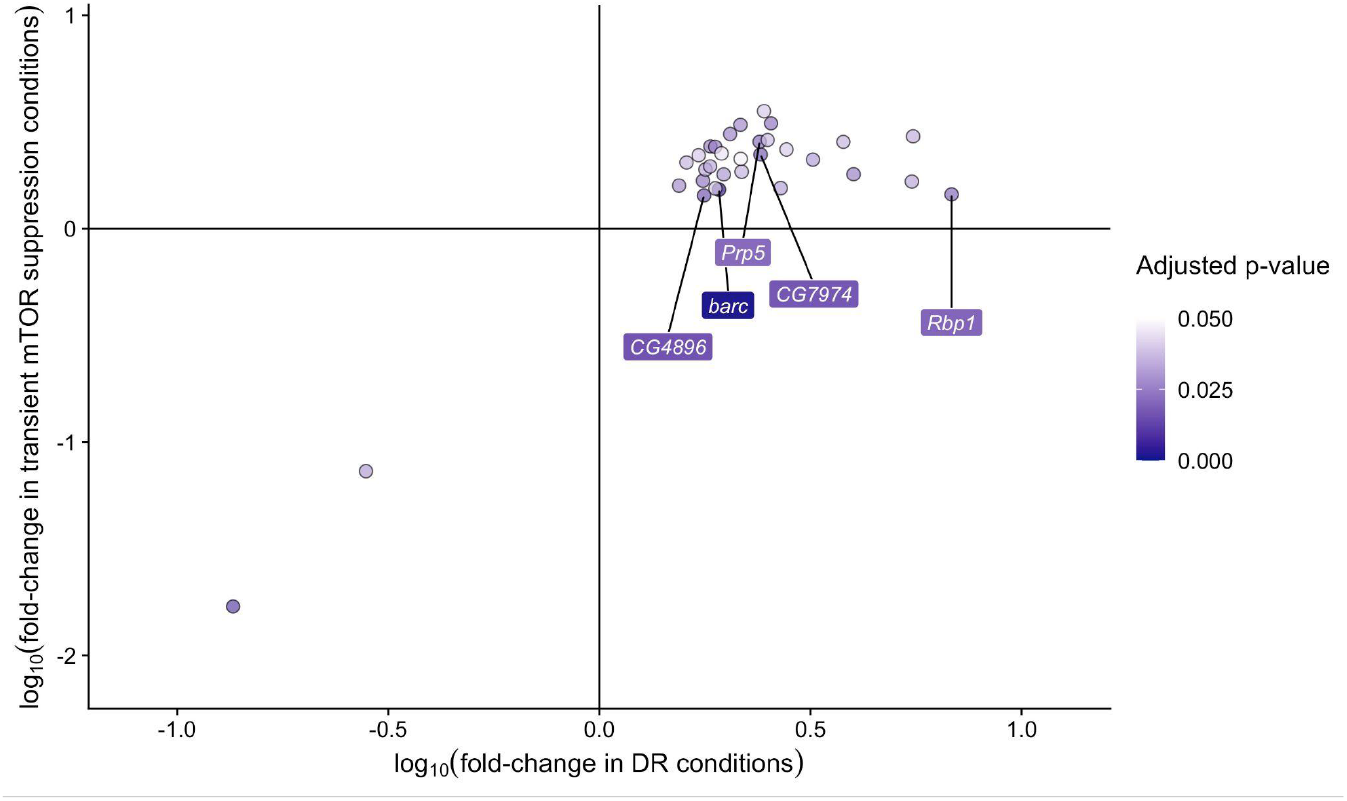
The spliceosome is modulated in both DR and transient mTOR suppressed conditions. Spliceosome genes which respond similarly between DR and transient mTOR suppression. 32 spliceosome-annotated genes were significantly differentially expressed (adjusted p-values ≤ 0.05) and showed concordant transcriptional change in response to transient mTOR suppression and DR in *Drosophila melanogaster*. The vast majority of these (30 of the 32) were upregulated. The five genes from this set we subsequently tested for lifespan phenotypes are indicated, and represent the top 5 significant genes in the DR treatment that also responded to mTOR suppression.

### Individual spliceosome components modulates lifespan

To test whether the spliceosome components associated with both pro-longevity treatments were able to modulate lifespan we knocked them down in adults using *in vivo* RNAi on rich diets (Gautrey et al., 2025). We tested the top 5 significant genes within the DR dataset that were also differentially expressed in the mTOR dataset, as the DR dataset overall appeared less sensitive or impactful on the spliceosome. We also only included those spliceosome components that went up in expression in response to the pro-longevity treatments. We further included *Sf3b1* as it is a known important spliceosome component and has previously been investigated in relation to cancer proliferation and mTOR signalling (Fuentes-Fayos et al., 2022; Han et al., 2022). Although knockdown of *CG4896* and *CG7974* did not affect survival, knockdown of *barc, Prp5* and *Sf3b1* each significantly reduced lifespan, whilst *Rbp1* knockdown significantly extended lifespan (Figure 2 and Table 3).

**Figure 2.**
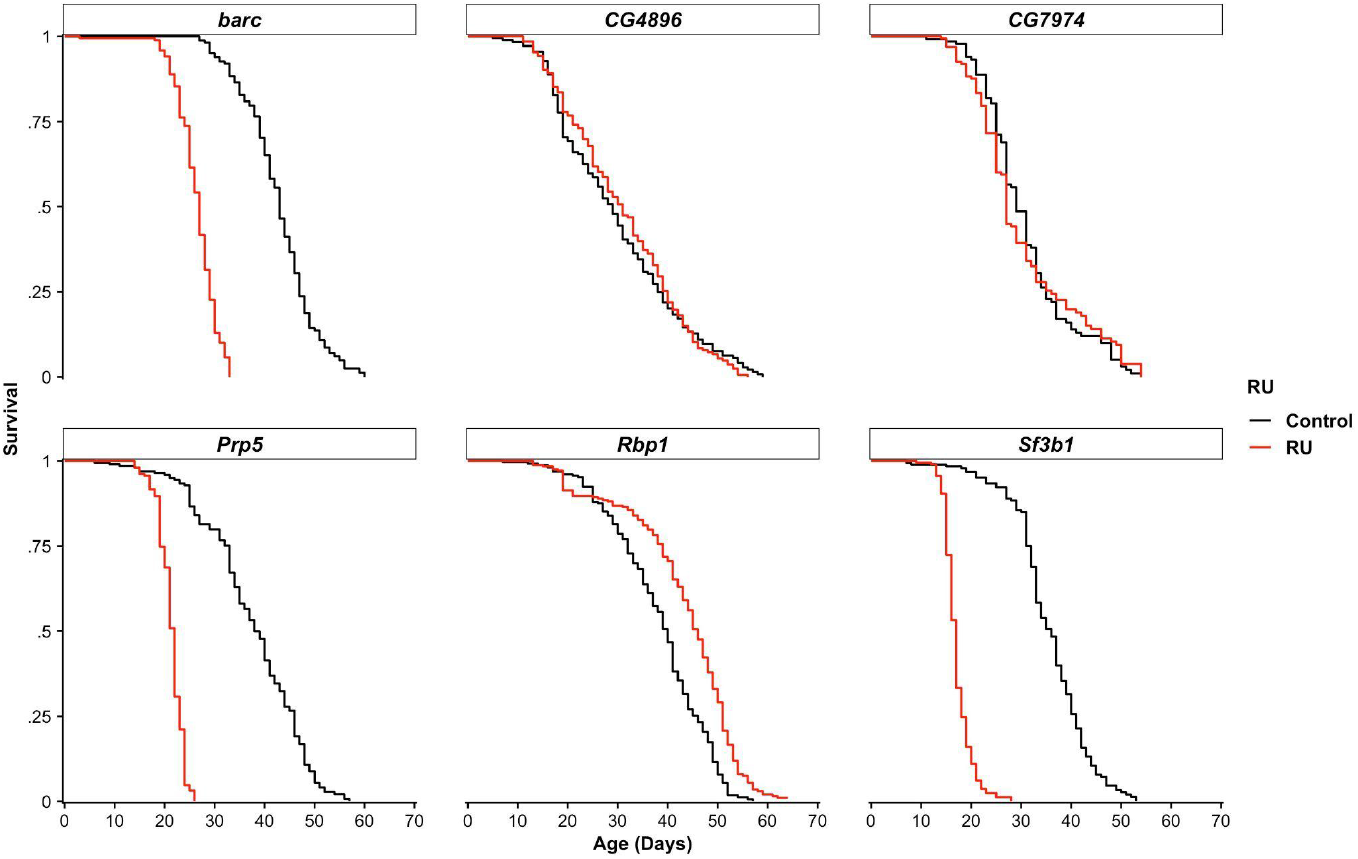
Knockdown of spliceosome genes modulates lifespan. Driving spliceosome gene RNAi with the *daughterless*-*GeneSwitch* (*daGS*) global conditional driver from age 4 days onwards reveals that different spliceosome components modulate lifespan in different directions in females (N ≥ 173 flies per condition). GeneSwitch activity is initiated by supplementation with RU486 (RU). Hazard ratios and p-values for the experiment are shown in Table 1.

### Knockdown of *Rbp1* reduces mortality when initiated later in adult life

Considering that conditionally driving *Rbp1* RNAi from early adult life extended lifespan, we wanted to test whether this treatment could also extend lifespan when activated in later life (Vaupel et al., 2003). We therefore repeated the *Rbp1* knockdown experiment, but with two additional conditions; *Rbp1* knockdown from 15 days onwards and *Rbp1* knockdown from 25 days onwards. Both of these treatments reduced mortality and thereby improved life expectancy (Figure 3 and Table 2). Knockdown of *Rbp1* can therefore improve life expectancy of flies irrespective of when the knockdown occurs.

**Table 1.** Statistics for knockdown of spliceosome genes. Data was analysed using cox proportional hazard mixed effects models using right-censoring where applicable (Therneau et al., 2003; Therneau, 2015), incorporating cage as a random effect and experimental batch as fixed effect, correcting for shared environmental effects from housing and growing conditions. Negative log hazard ratios indicate an increase in lifespan.

| Gene RNAi | $\log_e(\text{hazard ratio})$ | Standard error | N | P-value |
| --- | --- | --- | --- | --- |
| *barc* | 3.90 | 0.33 | 340 | < 0.0001 |
| *CG4896* | -0.14 | 0.11 | 375 | 0.20 |
| *CG7974* | 0.24 | 0.23 | 296 | 0.30 |
| *Prp5* | 3.44 | 0.24 | 404 | < 0.0001 |
| *Rbp1* | -0.69 | 0.13 | 501 | < 0.0001 |
| *Sf3b1* | 4.35 | 0.33 | 388 | < 0.0001 |

**Table 2.** Statistics for *Rbp1* knockdown at different timepoints, or overall. Data was analysed using coxme interval-based models, incorporating the cage ID as a random effect and experimental batch to capture shared environmental effects. As treatment was initiated later we coded this as a time-dependent covariate in the coxme models (McCracken et al., 2020; Therneau & Atkinson, 2016).

| Comparison | log<sub>e</sub>(hazard ratio) | Standard error | N | P-value |
| --- | --- | --- | --- | --- |
| Overall RU vs Control | -0.23 | 0.06 | 2800 | 0.00012 |
| RU at 15 days vs Control | -0.30 | 0.08 | 1525 | 0.00017 |
| RU at 25 days vs Control | -0.23 | 0.09 | 1547 | 0.00590 |

**Figure 3.**
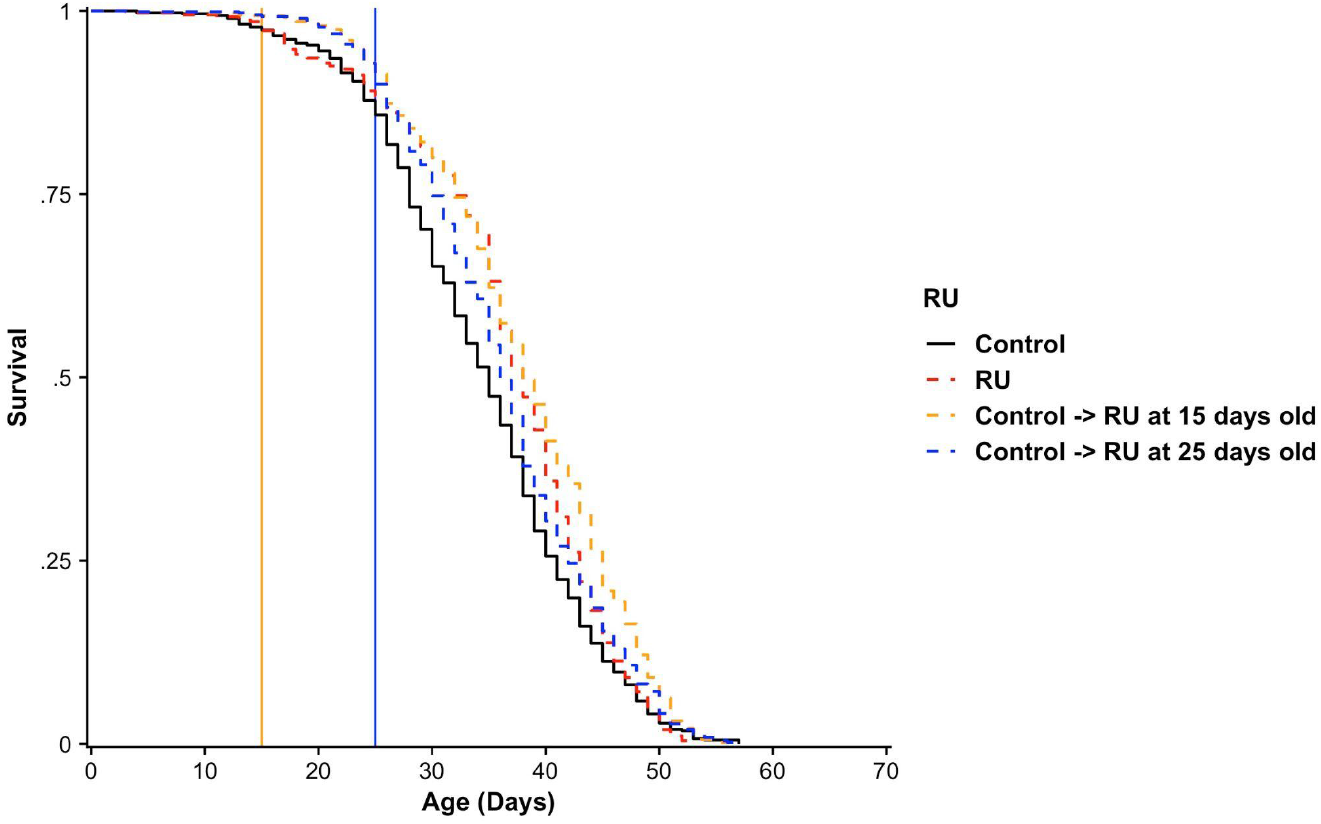
Knockdown of *Rbp1* reduces mortality risk irrespective of when it is triggered. Conditional activation of *Rbp1* RNAi reduced mortality irrespective of whether it was induced 4 days post-eclosion (as in Figure 2), or at later timepoints; 15 days post-eclosion or 25 days post-eclosion. *Rbp1* knockdown reduced mortality even when initiated at later timepoints (N ≥ 700 flies per condition, coxme interval-based model). The vertical yellow and blue lines signify the timepoints of RU supplementation.

## Discussion

Individual spliceosome components when experimentally reduced negatively affected lifespan but surprisingly *Rbp1* when knocked down increased lifespan. The direction of this effect is surprising as both pro-longevity treatments increase *Rbp1* expression. *Rbp1* knockdown alone is able to increase lifespan while in the context of pro-longevity treatments its increased expression is correlated with increased lifespan. This contradiction could be explained by interactions within the spliceosome or, alternatively, the spliceosome response observed is a compensatory response and it is actually the reduction of its downstream effects, possibly translation to protein that extends lifespan. We recently observed similar compensatory effects for a range of DR-responsive genes where similarly genes that increased in expression, when knocked down, increased lifespan (Gautrey et al., 2025).

In vertebrates, inhibition of many spliceosome genes severely affect many essential cellular processes and mutations in spliceosome genes are frequently observed in cancers (Olthof et al., 2022; Taylor & Lee, 2019). To a certain extent it was therefore unsurprising that we observed increased mortality upon conditional knockdown of *barc, Prp5* and *Sf3b1* in flies. However, unexpectedly the conditional knockdown of the SR protein-encoding gene *Rbp1* extended lifespan. In theory, knockdown of *Rbp1* should reduce spliceosome assembly at *Rbp1* binding sites in specific exonic splicing enhancers (ESEs), which therefore should negatively impact splicing fidelity of a subset of mRNAs. The mechanistic reason for this positive effect of knockdown of *Rbp1* on lifespan is at present unclear. However, interestingly, the human orthologue of *Rbp1, SRSF1* (Kim et al., 1992) has been previously suggested to constitute a rejuvenation factor (Plesa et al., 2023).

The spliceosome components which rapidly increased mortality risk when knocked down in this study are all recruited to the spliceosome at a later stage than the point at which Rbp1 plays a role; barc, Sf3b1 and Prp5 are all recruited at the pre-spliceosome stage (also known as Spliceosomal Complex A). Prp5 is recruited in order to bridge the U1 and U2 snRNPs together at the pre-mRNA (Liang & Cheng, 2015; Xu et al., 2004), whilst barc and Sf3b1 are recruited as part of the U2 snRNP (Abramczuk et al., 2017; van der Feltz & Hoskins, 2019). In contrast, *Rbp1* encodes an SR protein, a class of proteins which bind pre-mRNA and recruit early spliceosome components to splice sites even before the commitment complex (Spliceosomal Complex E) has formed (Kim et al. 1992; Jeong 2017). Therefore, perhaps a reduction (but not complete ablation) of spliceosome assembly, such as that which results from *Rbp1* suppression, benefits organismal health and longevity. One potential mechanism could be improved proteostasis by causing overall reduced protein synthesis (Basisty et al., 2018; Hipkiss, 2007). Perhaps suppression of components beyond recruitment towards the spliceosome causes the spliceosome to stall fully leading to a complete failure to effectively translate protein and thus lead to a severely truncated lifespan.

A second important result from this study is that *Rbp1* knockdown extends lifespan also when instigated later in adult life. Geroscience-based treatments that do not require lifelong treatment will be far easier to translate to the clinic (Kirkland, 2016). So far only mTOR suppression may be capable of achieving this across organisms (Bitto et al., 2016; Garratt et al., 2016; Simons et al., 2019). Rbp1 is especially interesting as it was also identified independently using a cell-based screen on reprogramming with subsequently beneficial phenotypes including in mice (Plesa et al., 2023). More generally, individual spliceosome components, rather than the spliceosome as a whole, may prove powerful targets for healthspan-modifying drugs.

## Methods

Fly media was composed of the following components, as previously described (Hayman et al., 2025): 6% cornmeal, 13% table sugar, 1% agar, 0.225% nipagin and 8% yeast (all w/v), with the addition of 0.4% (w/v) propanoic acid (Sigma-Aldrich) for fly growing bottles only. All experiments utilised the near globally-expressed *daughterless*-*GeneSwitch* (daGS) driver (Tricoire et al., 2009) to conditionally drive transgenes *in vivo*, thereby removing the potential effect of background genotype (Hayman et al., 2025). In experimental conditions requiring activation of daGS, RU486 (200 μM; Thermo Fisher Scientific) was supplemented, dissolved in ethanol and controls treatments received the same amount of ethanol. Media was split from a same batch to ensure the exact same food was used for both treatments. Previous experiments in our laboratory have found no effect of RU supplementation on lifespan using this same driver line (Gautrey et al. 2025; Hayman et al. 2025). All flies were maintained and grown at 25°C. The fly lines used are shown in Table S3. For survival experiments, flies were kept for mating for 2 days after eclosion, before being sorted under light carbon dioxide anesthesia, with females put into cages to assess survival as we described previously (Gautrey et al., 2025; Hayman et al., 2025; McCracken et al., 2020; Phillips & Simons, 2024).

## Acknowledgements

MJPS is a Sir Henry Dale fellow (Wellcome and Royal Society: 216405/Z/19/Z). Dan Hayman is a Vivensa Foundation ECR Fellow (ECRF24\13).

## Supplementary material

**Table S1.** Spliceosome-related Gene Ontology terms used to filter shared genes changing in mTOR suppressed and DR conditions.

|  |
| --- |
| Generation of catalytic spliceosome for first transesterification step |
| Generation of catalytic spliceosome for second transesterification step |
| Trans assembly of SL-containing precatalytic spliceosome |
| Cis assembly of pre-catalytic spliceosome |
| mRNA trans splicing, via spliceosome |
| Alternative mRNA splicing, via spliceosome |
| Regulation of alternative mRNA splicing, via spliceosome |
| Spliceosome conformational change to release U4 (or U4atac) and U1 (or U11) |
| mRNA splicing, via spliceosome |
| mRNA cis splicing, via spliceosome |
| Regulation of mRNA splicing, via spliceosome |
| Negative regulation of mRNA splicing, via spliceosome |
| Positive regulation of mRNA splicing, via spliceosome |
| U2-type prespliceosome |
| U2-type precatalytic spliceosome |
| U2-type catalytic step 1 spliceosome |
| U2-type catalytic step 2 spliceosome |
| Prespliceosome |
| Precatalytic spliceosome |
| Catalytic step 1 spliceosome |
| Catalytic step 2 spliceosome |
| U12-type prespliceosome |
| U12-type precatalytic spliceosome |
| U12-type catalytic step 1 spliceosome |
| U12-type catalytic step 2 spliceosome |
| Spliceosome-depend formation of circular RNA |
| U2-type prespliceosome assembly |
| Regulation of mRNA cis splicing, via spliceosome |
| Negative regulation of mRNA cis splicing, via spliceosome |
| Positive regulation of mRNA cis splicing, via spliceosome |

**Table S2.**
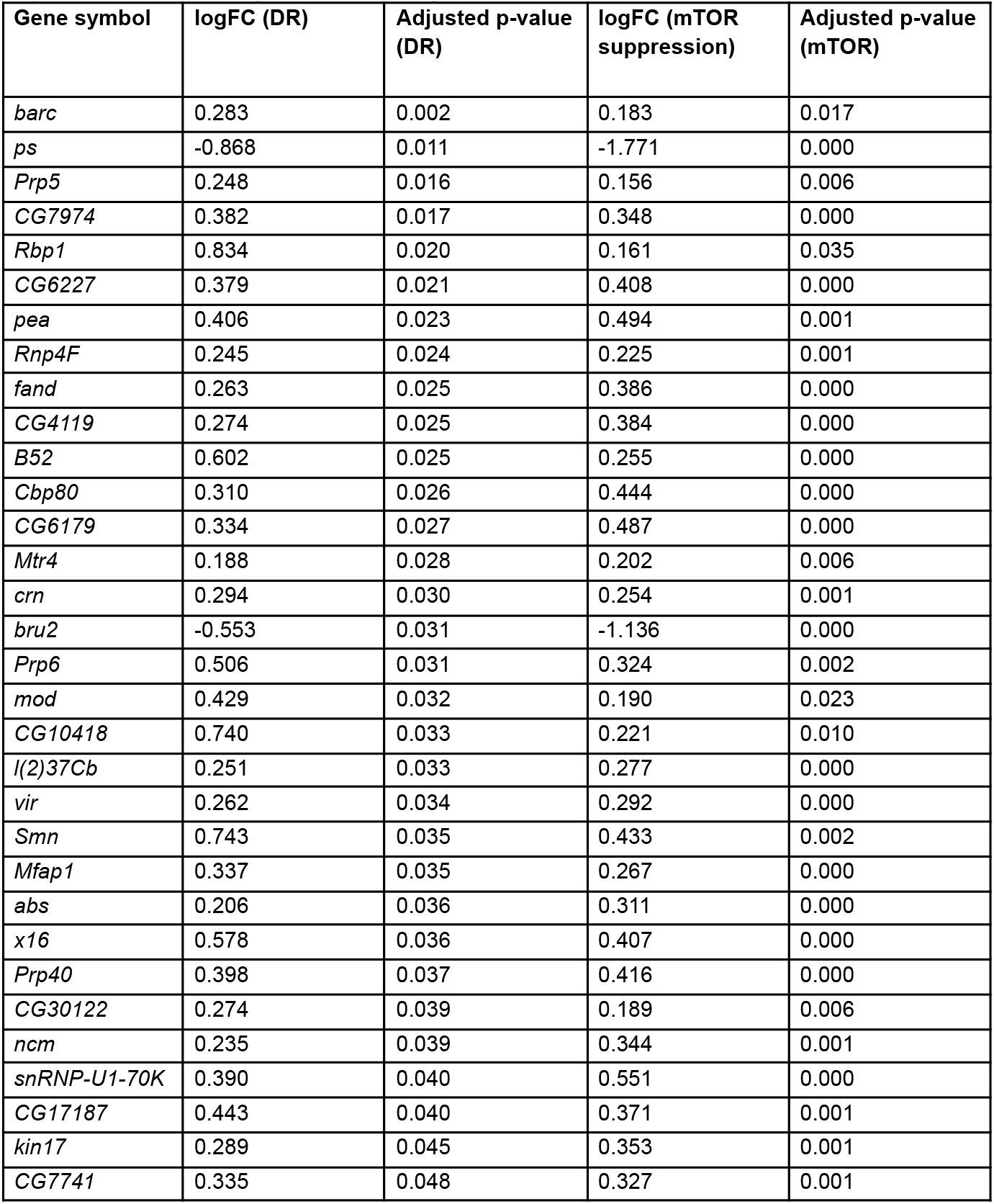
Shared dysregulated spliceosome genes between mTOR suppression and DR. 30 spliceosome-annotated genes are upregulated upon both transient mTOR suppression and DR, whilst 2 spliceosome-annotated genes are downregulated in both treatments.

**Table S3.**
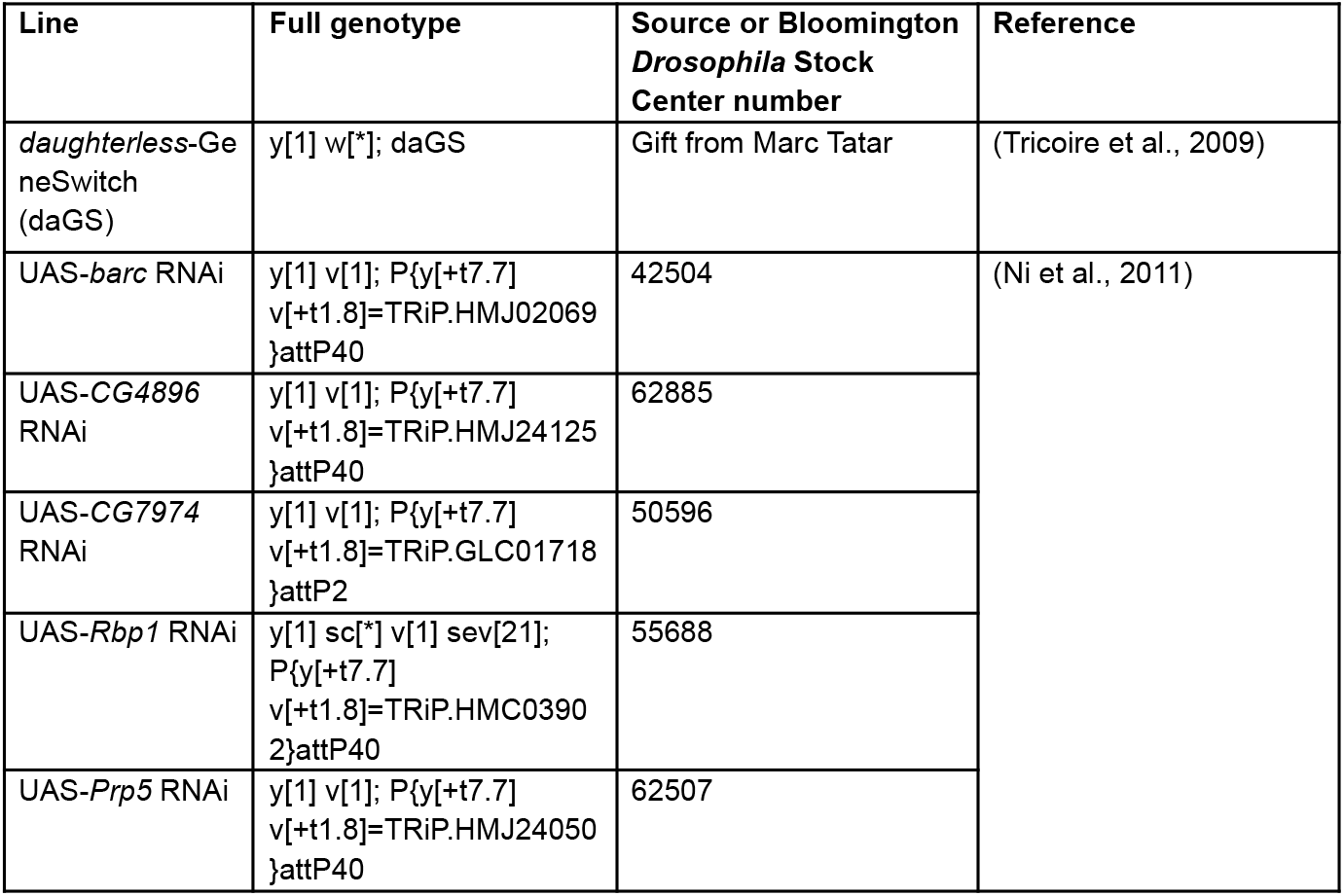
Fly lines used in this manuscript, with source stock and/or publication.

